# MICS1 is the Ca^2+^/H^+^ antiporter of mammalian mitochondria

**DOI:** 10.1101/2021.11.11.468204

**Authors:** Shane Austin, Ronald Mekis, Sami E. M. Mohammed, Mariafrancesca Scalise, Christina Pfeiffer, Michele Galluccio, Tamara Borovec, Katja Parapatics, Dijana Vitko, Nora Dinhopl, Keiryn L. Bennett, Cesare Indiveri, Karin Nowikovsky

## Abstract

Mitochondrial Ca^2+^ ions are crucial regulators of bioenergetics, cell death pathways and cytosolic Ca^2+^ homeostasis. Mitochondrial Ca^2+^ content strictly depends on Ca^2+^ transporters. In recent decades, the major players responsible for mitochondrial Ca^2+^ uptake and release have been identified, except the mitochondrial Ca^2+^/H^+^ exchanger (CHE). Originally identified as the mitochondrial K^+^/H^+^ exchanger, LETM1 was also considered as a candidate for the mitochondrial CHE. Defining the mitochondrial interactome of LETM1, we identified MICS1, the only mitochondrial member of the TMBIM family. Applying cell-based and cell-free biochemical assays, here we demonstrate that MICS1 is responsible for the Na^+^- and permeability transition pore-independent mitochondrial Ca^2+^ release and identify MICS1 as the long-sought mitochondrial CHE. This finding provides the final piece of the puzzle of mitochondrial Ca^2+^ transporters and opens the door to exploring its importance in health and disease, and to developing drugs modulating Ca^2+^ exchange.

## Introduction

Ion homeostasis is important for maintaining mitochondrial function. The dynamic balance of ions to maintain function is achieved by various cycles, which facilitate the interplay of cations, via the K^+^, Na^+^, and Ca^2+^ cycles. Loss of this balance leads to several consequences in the organelle and ultimately the cell. These include mitochondrial swelling, disrupted cristae structure, deregulated bioenergetics and may result in cell death. Intracellularly, mitochondria have been established as major sinks of Ca^2+^, an ion of comparatively low concentration to K^+^ and Na^+^. The role of mitochondrial Ca^2+^ buffering has been extensively studied (Giorgi et al., 2018; Pallafacchina et al., 2018), yet some of the players in maintaining this Ca^2+^ balance have not been identified (De Stefani et al., 2016; Urbani et al., 2020). One of the missing pieces in this molecular puzzle is the Na^+^-independent Ca^2+^ efflux pathway, a putative Ca^2+^/H^+^ exchanger (CHE). This exchanger, whose existence has been postulated since the 1970s (Carafoli et al., 1974) is critical for maintaining mitochondrial Ca^2+^ levels and plays an important role in mitochondrial functions.

To date, several studies have investigated the molecular identity of mitochondrial CHE, one of the likely candidates being LETM1. LETM1 first attracted interest when it was found to be associated with seizures in the Wolf Hirschhorn syndrome (Endele et al., 1999). Over the years, numerous studies characterized LETM1, a single transmembrane domain-containing protein, as the mitochondrial K^+^/H^+^ exchanger (KHE) (Hasegawa and van der Bliek, 2007; Hashimi et al., 2013; McQuibban et al., 2010; Nowikovsky et al., 2004; Nowikovsky et al., 2007). The proposal that LETM1 could be the CHE was based on a *Drosophila* S2 genome-wide RNAi screen of modulators of mitochondrial Ca^2+^ transport (Jiang et al., 2009). Subsequent studies have confirmed an involvement of LETM1 in Ca^2+^ and K^+^ transport but key questions remained (Austin and Nowikovsky, 2019, 2021; Nowikovsky and Bernardi, 2014). Perhaps the most important is how can a single transmembrane protein mediate a process of ion exchange, and it appeared possible that LETM1 could fulfill its function(s) as a multimer, or as part of a protein complex. The first possibility was addressed by Shao et al., who presented cryo EM structures of LETM1 oligomers, which facilitated pH-dependent movement of Ca^2+^ in a cell-free system (Shao et al., 2016). Whether LETM1 is part of a protein complex remains unaddressed.

In this study, we searched for partners of LETM1 and found the interactor mitochondrial Morphology and Cristae Structure 1 (MICS1), a member of the TMBIM family, which has been implicated in the regulation of intracellular Ca^2+^ by a number of studies (Carrara et al., 2012; Hung et al., 2011; Kim et al., 2021; Lisak et al., 2015; Liu, 2017; Rojas-Rivera and Hetz, 2015). Interestingly, MICS1 is the only species with a mitochondrial localization (Oka et al., 2008) while other TMBIM family members are localized to the ER, Golgi and the plasma membrane (Rojas-Rivera and Hetz, 2015). Functional characterization of the TMBIM family members has been generally addressed with mammalian cell culture and animal models, especially investigating their role in Ca^2+^ regulation. In fact, MICS1 was identified as a regulator of calcium and apoptosis (Lisak et al., 2015; Oka et al., 2008). Here, we demonstrate that MICS1 is the long-sought mitochondrial CHE, a crucial component of mitochondrial Ca^2+^ homeostasis.

## Results

### MICS1 interacts with LETM1

We determined the interactome of LETM1 using affinity purification mass spectrometry (AP-MS) from whole cell lysates and isolated mitochondria (**Figure 1A**). As few studies have used the limited amounts of material from isolated organelles for AP-MS, we assessed the suitability of our modified method to investigate organellar interactomes (**Figure 1–figure supplement 1A**). Using the mitochondrial Ca^2+^ uniporter (MCU) as a benchmark inner mitochondrial membrane protein, we confirmed the ability of our method to detect the members of the published core interactome except for the tertiary interactor MICU2, which interacts with MICU1 (Sancak et al., 2013) (**Figure 1–figure supplement 1B**). Thus, the method was sufficiently robust to cover approximately 75% of a mitochondrial core interactome (**Figure 1–figure supplement 1C**). We then determined the LETM1 interactome. Data obtained from both the whole cell and isolated mitochondria data sets were reproducible with an overlap of 31 proteins that interacted in both approaches (**Figure 1–figure supplement 1D**) including TBK1, a protein previously observed to interact with LETM1 in similar AP-MS studies (Li et al., 2011).

**Figure 1.**
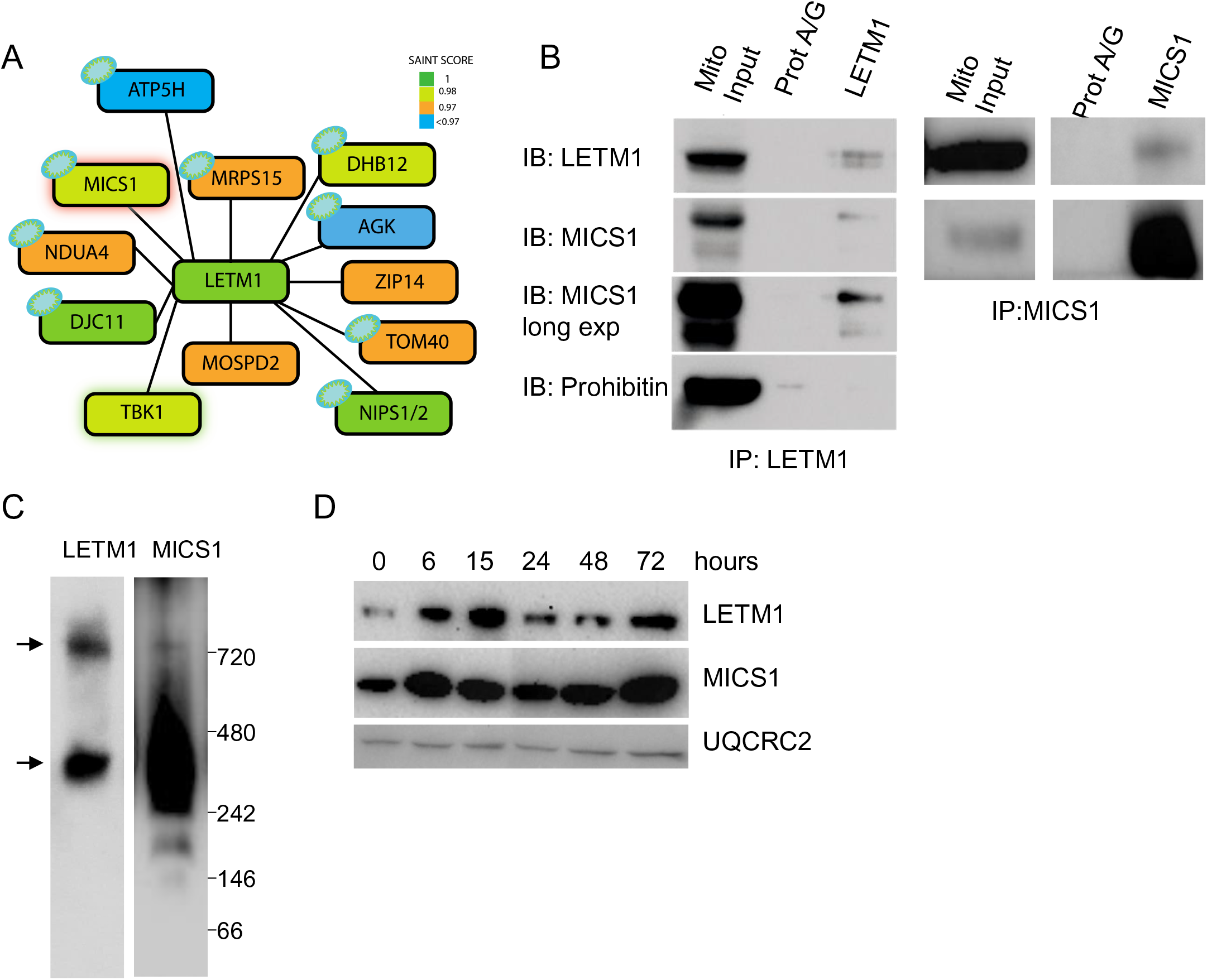
LETM1 and MICS1 interact. **(A)** LETM1 interactome as determined by affinity purification mass spectrometry (AP-MS). All high confidence interaction partners of LETM1 are shown as nodes. Node color indicates SAINT score, a probability-based measure of interaction confidence. Data are from a single MS experiment. See also **Figure1–figure supplement 1 (B)** Co-immunoprecipitation of MICS1 and LETM1 protein in tandem. Mitochondria were crudely isolated from HEK293 cells and used for immunoprecipitation. The input represents the mitochondrial crude lysate used as input for the co-IP, LETM1 was immunoprecipitated (left panel, IP:LETM1) using a LETM1 polyclonal antibody and Protein A/G agarose beads (ProtA/G). ProtA/G beads alone were used as a negative control for binding, immunoprecipitates were immunoblotted (IB) for the indicated proteins to demonstrate interaction. Prohibitin was used as a control to illustrate no nonspecific binding of inner mitochondrial membrane proteins complexes. The right panel illustrates the converse experiment, MICS1 was precipitated (right panel, IP: MICS1) using a MICS1 polyclonal antibody. **(C)** Native immunoblot of LETM1 (left) and MICS1 (right) show that both proteins can be found in protein complexes of the same size (arrows), MICS1 additionally resides in other protein complexes. **(D)** Immunoblot analysis of LETM1 and MICS1 expression on conditions of reduced FBS (0.5%) in culture media. Mitochondrial complex III integral subunit UQCRC2 is used as loading control.

One protein that was immediately of interest was MICS1, which is an inner mitochondrial membrane protein with 6-8 transmembrane domains depending on the prediction tool used. Similar to LETM1, MICS1 has already been shown to be involved in the regulation of mitochondrial structure (Oka et al., 2008; Seitaj et al., 2020).

Specifically relying on crudely isolated mitochondria to perform co-immunoprecipitation experiments, we were able to confirm that LETM1 does indeed interact with MICS1 (**Figure 1B**). Probing for mitochondrial Prohibitin demonstrated this was not an enrichment of membrane-associated proteins, but rather a specific complex (**Figure 1B-left**). The same result was obtained when MICS1 was immune-precipitated, with LETM1 being present in the same complex (**Figure 1B-right**). LETM1 forms high molecular weight complexes migrating at approximatively 400 and 720 kDa. Blue native gel electrophoresis (BNGE) detected LETM1 and MICS1 in same complexes of 720 kDa and ∼ 400 KDa (**Figure 1 C**). Applying low serum conditions to enhance MICS1 levels (Oka et al., 2008), the alterations of MICS1 amounts were paralleled by LETM1, while in contrast the amounts of the mitochondrial protein UQCRC2 did not change (**Figure 1D**).

### MICS1 depletion impairs mitochondrial bioenergetics and morphology

Functional characterization of MICS1 being limited, we first generated MICS1 stable knockdown by short hairpins targeting various exons. Stable knockdown cells had up to 80 % reduced MICS1 levels compared to scrambled controls and were accompanied by a proportional decrease of LETM1 (**Figure 2A**). The proliferation rate of MICS1KD1 in glucose-containing media was modestly reduced compared to controls, with only the final time point being significantly affected (**Figure 2B**). While no significant difference in any respiratory parameter was observed with glucose as the substrate (**Figure 2C-D**), galactose-dependent respiration was severely compromised in MICS1KD cells (**Figure 2E-F**), indicating that MICS1 impacts on mitochondrial function. To further address the specific function of MICS1 in mitochondrial morphology and cation homeostasis, we generated MICS1 knockout HEK293 and HeLa cells by CRISPR/Cas9 genome editing. At the gene expression level, we obtained HEK293 and HeLa knockout individual clones that entirely abrogated the transcript levels of MICS1. At the protein level, the total knockout was confirmed in HeLa cells clone IIIF3 (HeLa MICS1KO) and in HEK293 cells clone IIF1 (HEK293 MICS1KO#1). In several other clones, translation was not entirely abolished, like in HEK293 clone IE12 (HEK293 MICS1KO#2) (**Figure 3A and E**). Comparison of HEK293 cell proliferation rates indicated that while the complete loss of MICS1 did not affect cell growth, growth of mutant cells with residual MICS1 expression was significantly slowed (**Figure 3B**), similarly to MICS1KD (**Figure 2B**), suggesting a potential cellular adaptation in a full KO.

**Figure 2.**
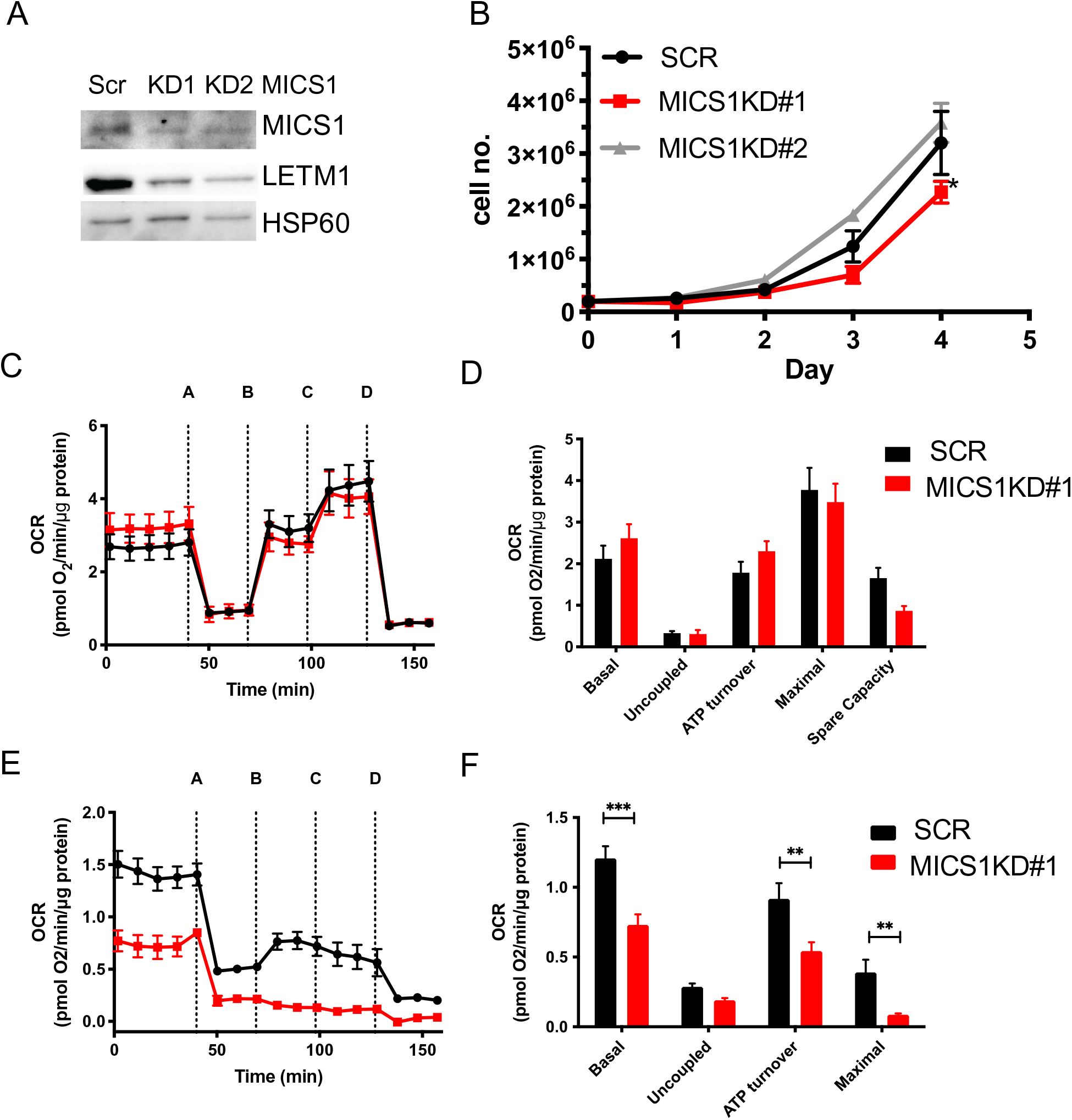
MICS1KD decreases LETM1 and mitochondrial bioenergetics. **(A)** Western blot analysis of LETM1 and MICS1 in HEK293 MICS1WT cell with scramble shRNA (scr) or two different MICS1 knockdowns (KD#1 or KD#2). HSP60 served as a loading control. **(B)** Proliferation curve of MICS1WT (WT) with a scrambled construct compared to MICS1KD cells (KD) over 4 days using a trypan blue exclusion assay to count cells. Data are means ± SEM (n=3), at 96h statistical analysis using an unpaired student’s t-test (***p<0.001). **(C-F)** Cellular bioenergetics of MICS1KD cells in various nutrient conditions. Oxygen consumption rate of WT cells with a scrambled control (“WT”) and MICS1KD#1 cells grown in **(C)** 25 mM glucose, **(E)** 10 mM galactose for 24 hours before measurement. Data are representative of at least 3 independent experiments. Shown are mean data of triplicate measurements ± SEM. Inhibitors as indicated: A-oligomycin (0.5 μM), B & C-FCCP (0.2 μM each), D-antimycin A/rotenone (0.5 μM). **(D & F)** Bar charts of XF experiment traces (C & E), data are means of multiple time points after experiment start or drug addition of at least three independent experiments ± SEM. (n=3). Statistical analysis using an unpaired student’s t-test (\*\**p*<0.01, \*\*\**p*<0.001).

**Figure 3.**
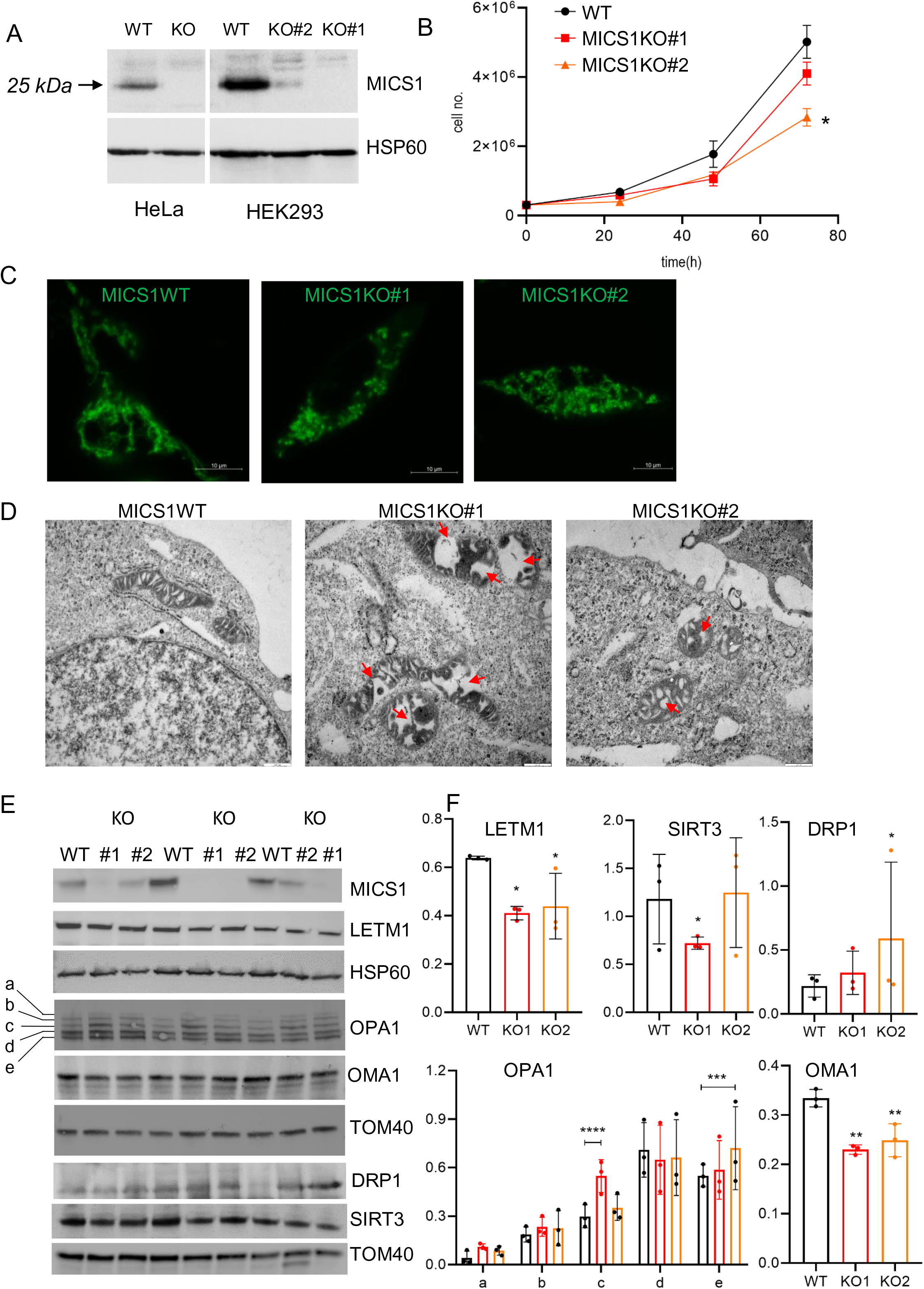
MICS1KO causes mitochondrial matrix swelling and cristae disorganization. **(A)** Western blot analysis of MICS1 in control and targeted HeLa and HEK293 clones, HSP60 served as loading control. **(B)** Proliferation assay of HEK293 cells in function of MICS1. Graph shows the mean of three individual counts, Two-way ANOVA with Dunnett’s multiple comparisons test performed against MICS1WT *p=0.0155. **(C)** Live imaging of HEK293 MICS1WT and KO cells stained with MitoTracker Green FM. Bars: 10 μm **(D)** Alteration of the mitochondrial ultrastructure shown by transmission electron microscopy, red arrow pointing to dilated matrix. Wider mitochondria in middle and right panel compared to controls, a middle panel showing the strongest phenotype of matrix width and cristae forms. **(E)** Isolated mitochondria from three independent replicates of HEK293 MICS1WT and MICS1KO#1 (#1) and KO#2 (#2) were analyzed by immunoblotting using the indicated antibodies, HSP60 and TOM40 served as mitochondrial loading controls. **(F)** Densitometric analysis of the bands in (E) normalized to loading control, bar graph of three individual counts, One-way ANOVA with Bonferroni’s multiple comparisons test performed against MICS1WT *p<0.05, **p<0.008, Two-way ANOVA with Bonferroni’s multiple comparisons test performed for the OPA1 statistics against MICS1WT, ***p=0.0009, ****p<0.0001.

We performed transmission electron microscopy to study mitochondrial ultrastructure of MICS1-deficient cells that had a null mutation or still had residual levels of the MICS1 protein. As previously shown for HeLa and HAP cells (Oka et al., 2008; Seitaj et al., 2020), compared to wild-type cells HEK293 MICS1KO#1 and MICS1KO#2 displayed fragmented and less elongated mitochondria, respectively (**Figure 3C**). Electron micrographs showed mitochondria with swollen sections and altered cristae structures when MICS1 was deleted, cristae being also affected in the incomplete MICS1KO (**Figure 3D, arrows**). Since OPA1 is known to control cristae volume and cristae junction organization, which is a crucial determinant for mitochondrial cytochrome c retention (Del Dotto et al., 2017; Olichon et al., 2003), we investigated whether cristae changes were associated with changes in the cleavage pattern of OPA1 isoforms. Increased c and e subunits were apparent in both complete and partial KO compared to controls (**Figure 3E-F**). OPA1 c and e forms are the cleavage products of OMA1. We found significantly reduced levels of OMA1, in line with the autocatalytic degradation of activated OMA1 (**Figure 3E-F**). In addition, DRP1 was upregulated (**Figure 3E-F**), supporting the shift of the mitochondrial morphology towards increased fission and consistent with stress-sensitive activation of OMA1 and OMA1-dependent OPA1 cleavage. Our western blot analysis further confirmed the depletion of MICS1 in MICS1KO#1 and strong reduction in MICS1KO#2 and the proportional decrease of LETM1 (**Figure 3 E-F**).

### Mitochondrial KHE requires LETM1 and MICS1

Based on the interaction of MICS1 with LETM1 and on the implication of LETM1 in mitochondrial K^+^/H^+^ exchange (Nowikovsky et al., 2012), we asked whether MICS1 contributes to KHE activity. Light scattering methods have been classically used to monitor the swelling of mitochondria (Bernardi, 1999; Mitchell, 1966). Using acetate-based cationic salts to measure the swelling of isolated mitochondria is a robust and accurate method to assess relative KHE activity in these organelles. HeLa and HEK293 MICS1KO mitochondria had a significantly reduced rate of swelling in potassium acetate buffer (**Figure 4 A-D**), indicating reduced KHE activity, as also seen for both K^+^ and Na^+^ salts in mitochondria from LETM1KD cells (**Figure 4E-F**), see also (Austin et al., 2017). Re-expression of MICS1 in HeLa MICS1KO cells restored swelling to wild-type levels (**Figure 4A-B**), confirming a correlation between MICS1 and KHE activity. Together with the proportional decrease of LETM1 in MICS1 knockdown or knockout (**Figure 2A**), these data suggested that MICS1 depletion may reduce the KHE activity by destabilizing LETM1. Importantly, these findings demonstrate that LETM1-mediated active KHE activity requires the presence of MICS1.

**Figure 4.**
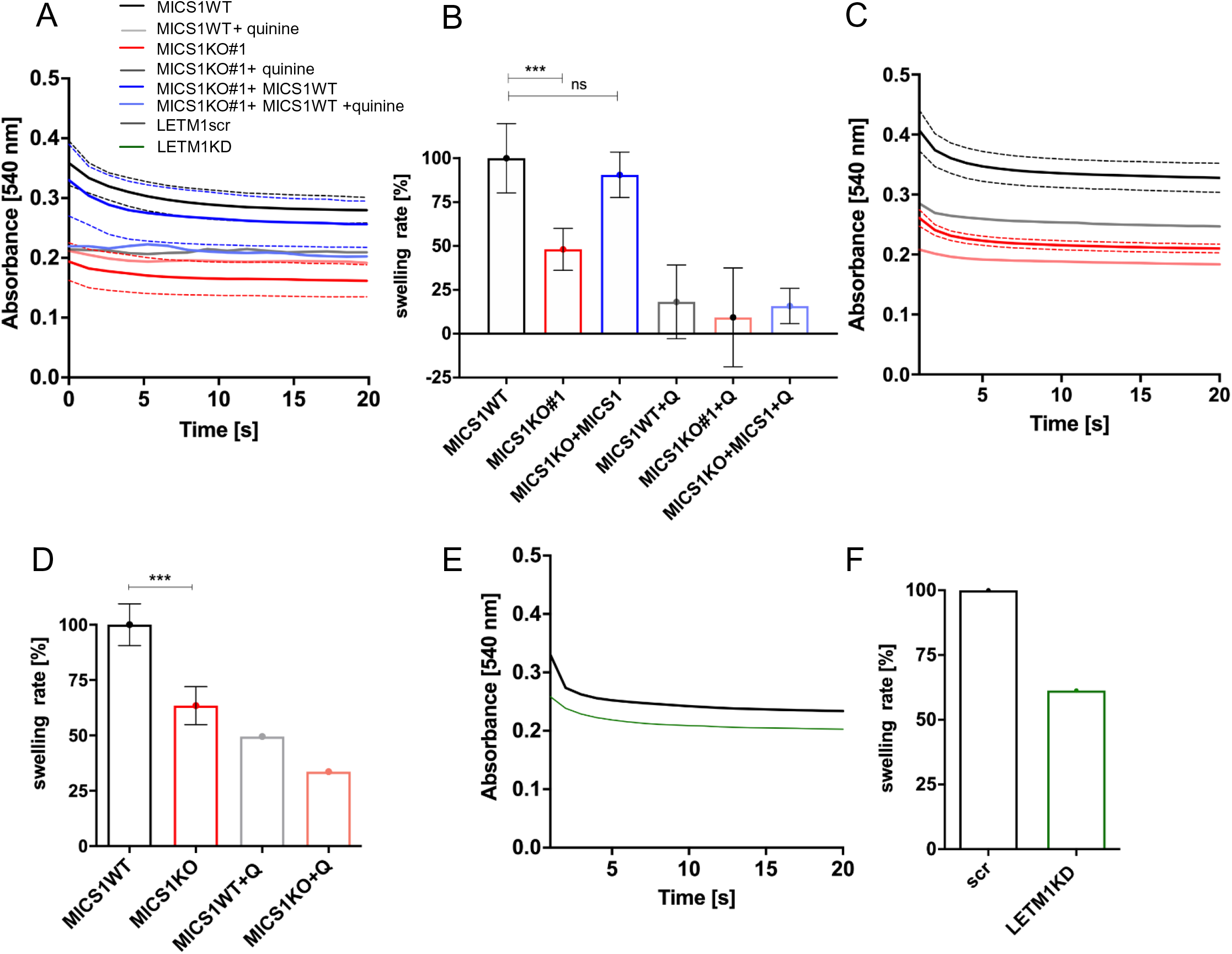
MICS1 and LETM1 are involved in mitochondrial KHE activity. KOAc-induced swelling was measured in mitochondria derived from HEK293 MICS1WT, MICS1KO and MICS1KO cells stably re-expressing MICS1WT **(A-B)**, HeLa MICS1WT and MICS1KO **(C-D)** and HeLa LETM1 scramble and LETM1 KD **(E-F)** cells. MICS1WT: black traces, MICS1KO: red traces, MICS1KO + MICS1WT: blue traces, LETM1scr: black trace, LETM1KD: green trace. **(B)** Quantification of swelling amplitudes from independent experiments (n=3) HEK293 MICS1WT (black bar, 100 ± 19.71) and HEK293 MICS1KO (red bar, 48.08 ± 11.906). Complementation of MICS1KO with re-expression of *MICS1WT* restored swelling rates (blue bar, 90.55 ± 12.93). **(D)** Similar differences in swelling capacities were obtained between HeLa MICS1WT (black bar; 100 ± 9.47) and HeLa MICS1KO (red bar; 63.48 ± 8.60). Lower basal optical density indicates swollen matrix prior KOAc addition; Inhibition of KHE with quinine in HEK293 cells: WT grey bar, 18.14 ± 21.02; KO#1 pink bar, 9.33 ± 28.17; MICS1KO+MICS1, 15.79 ± 10.04. Statistical analysis: One-Way ANOVA with Bonferroni correction (*p <0.05, **p <0.01, ***p <0.001). See also **Figure 4–figure supplement 1**.

### MICS1 mediates mitochondrial Na^+^-independent Ca^2+^ efflux

As the TMBIM protein family controls intracellular Ca^2+^ and previous work has proposed a Ca^2+^ channel function linked to pH sensitivity for the bacterial TMBIM homolog BsYetJ (Guo et al., 2019), we asked whether MICS1 controls mitochondrial Ca^2+^ homeostasis by mediating Ca^2+^/H^+^ exchange. To this end, we performed mitochondrial Ca^2+^ uptake and release assays in digitonin-permeabilized HEK293 cells pulsed with external Ca^2+^. To focus on H^+^-dependent Ca^2+^ fluxes and exclude Na^+^-dependent Ca^2+^ fluxes, we used the NCLX inhibitor CGP37157. MICS1WT and MICS1KO mitochondria exhibited similar rates of energy-dependent Ca^2+^ uptake (**Figure 5A and C and Figure 5–figure supplement 1**). The MCU inhibitor ruthenium red (RR) induced Ca^2+^ release from wild-type mitochondria, confirming that mitochondria can extrude matrix Ca^2+^ through an NCLX independent pathway, which is widely assumed to be a CHE. Residual matrix Ca^2+^ was then initiated by the pore-forming peptide alamethicin, or by FCCP, the protonophore that collapses the proton gradient. Wild-type mitochondria released Ca^2+^ to corresponding levels of total Ca^2+^ uptake (**Figure 5A**). Remarkably, MICS1KO mitochondria displayed decreased to absent RR-induced mitochondrial Ca^2+^ release, which was proportional to the depletion of MICS1 mitochondria (**Figure 5A-D** red and orange traces). Released total mitochondrial Ca^2+^ through alamethicin reached similarly high level in MICS1KO as in MICS1WT, confirming comparable levels of matrix Ca^2+^ (**Figure 5A**). Re-expression of MICS1 in HEK293 MICS1 KO was able to restore Ca^2+^ efflux (**Figure 5A**).

**Figure 5.**
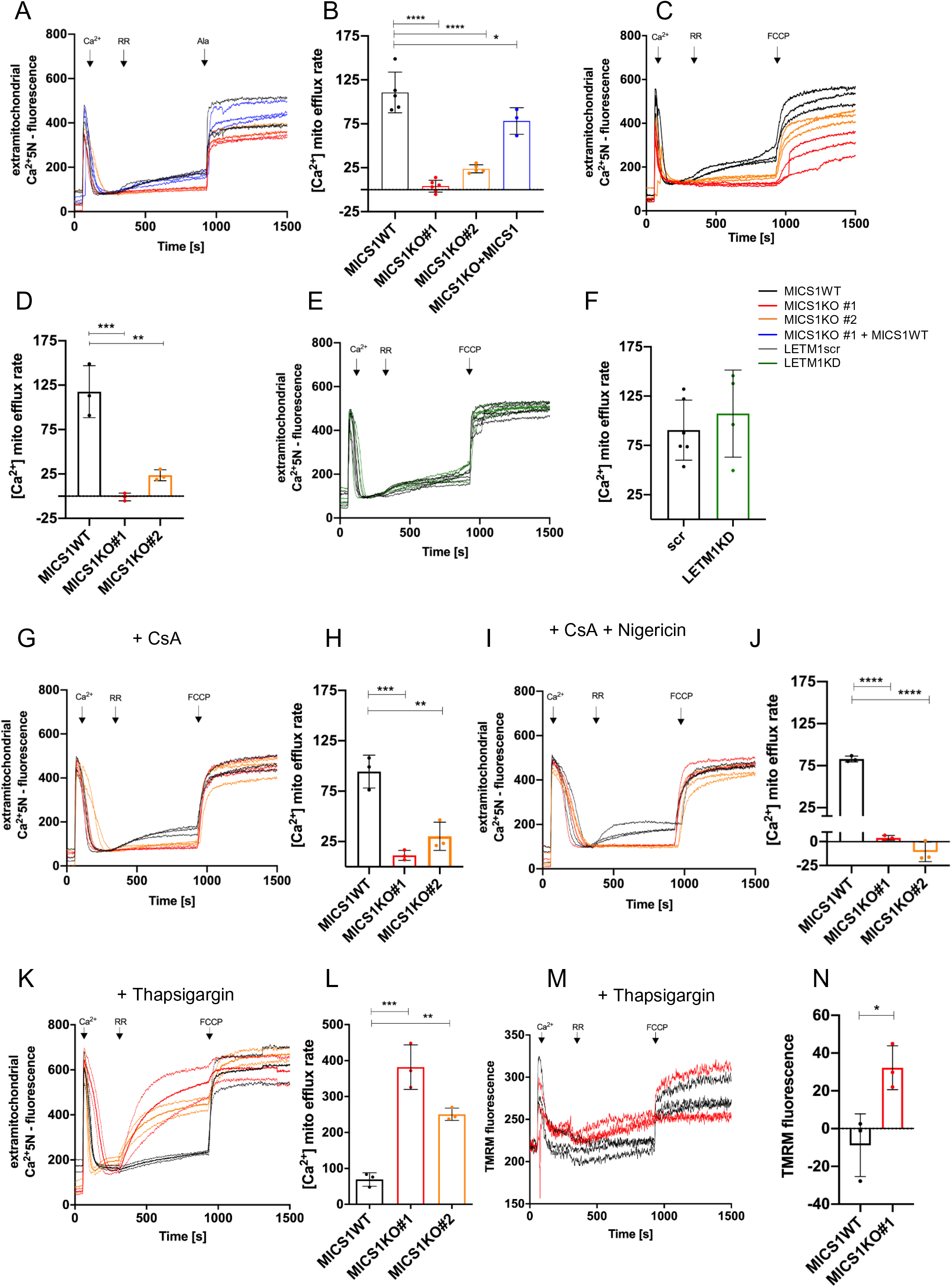
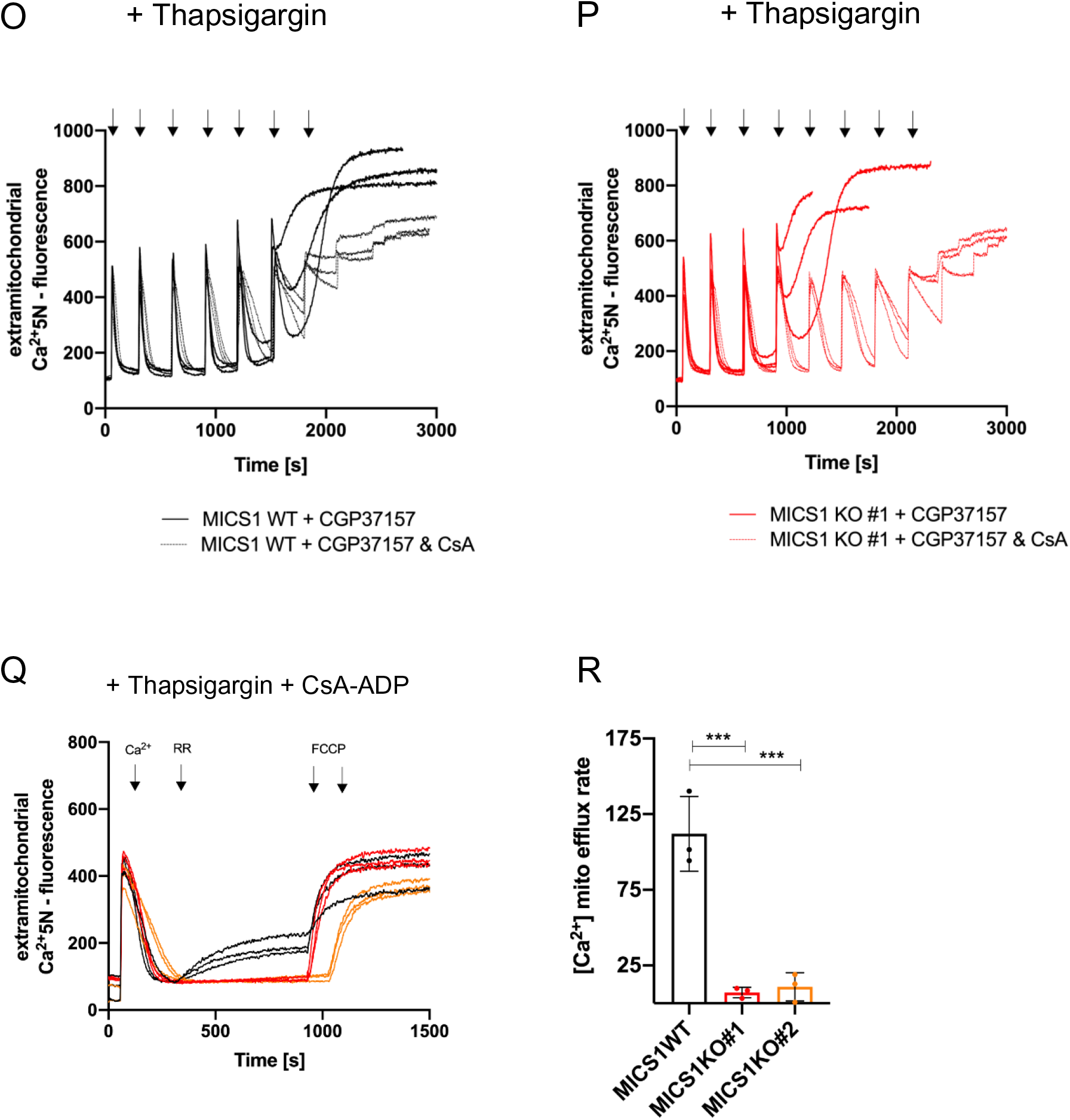
MICS1 controls Na^+^-independent Ca^2+^ release. Ca^2+^ uptake/release dynamics are shown as extramitochondrial Ca^2+^ changes of fluorescence intensities of Calcium Green 5N (Ca^2+^ 5N) (0.24 μM) **(A-L** and **P-Q)** and membrane potential as change of fluorescence intensities of TMRM (330 nM) **(M-N)** corresponding to the measurement of Ca^2+^ fluxes in **(K)**. Experiments were performed using permeabilized HEK293 MICS1WT, MICS1KO (KO#1, KO#2) (**A-D)** and **(G-R),** MICS1KO#1 + MICS1WT cells was included in **(A-B)**, and HEK293 LETM1 scr and LETM1KD cells **(E-F)** in presence of CGP37157 (2 μM). Ca^2+^ (10 μM), RR (0.2 μM) and FCCP (2 μM) or alamethicin (2.5 μM) were added when indicated. CsA was added two minutes before measurements in **(G-J** and **Q-R)**, nigericin was added 1 min before measurements in absence of thapsigargin or after Ca^2+^ uptake in thapsigargin experiments to prevent slowed Ca^2+^ uptake dynamics in **(I-J)**, thapsigargin (1 μM) was added as indicated in **K-R**, and ADP as indicated in **(Q-R)**. Quantification of Ca^2+^ release rates from independent experiments (n=3) (t: 300-920 s) and statistical analysis: One-Way ANOVA with Bonferroni correction (*p <0.05, **p <0.01, ***p <0.001, ****p <0.0001). See also **Figure 5–figure supplement 1** for quantification of Ca^2+^ uptake. Quantification of TMRM performed with unpaired two-sided t-test (Welsh correction), *p <0.05. **(O-P)** CRCs showing that absence of MICS1 supersensitizes mitochondria to Ca^2+^-induced PTP opening by thapsigargin. See also **Figure 5–figure supplement 2** for CRCs. In absence of thapsigargin. Permeabilized HEK293 MICS1WT **(O)** and MICS1KO#1 **(P)** cells exposed or not to CsA were subjected to sequential Ca^2+^ bolus of 5 µM Ca^2+^ and fluorescence intensity was recorded. **(Q-R)** Thapsigargin-dependent Ca^2+^ uptake/release experiments repeated in presence of CsA and ADP in MICS1WT and MICS1KO#1 and KO#2 showing the suppression of Ca^2+^ release, quantifications using One-Way ANOVA with Bonferroni correction.

Since MICS1 and LETM1 interact, and LETM1 was proposed as the mitochondrial CHE, we next sought to address once again Ca^2+^ fluxes in HEK293 LETM1 KD under the same conditions. The presence or absence of LETM1 (**Figure 4–figure supplement 1**) did not alter Ca^2+^ uptake (**Figure5–figure supplement 1**) nor the Na^+^-independent Ca^2+^ fluxes (**Figure 5E-F**). To assess whether the permeability transition pore (PTP) contributes to the recorded Ca^2+^ fluxes, we repeated Ca^2+^ uptake/efflux assays in presence of cyclosporin A (CsA), the PTP desensitizer (Basso et al., 2008). MICS1WT displayed comparable Ca^2+^ efflux as in the absence of CsA, confirming that Na^+^-independent Ca^2+^ release was also independent of PTP flickering or opening (**Figure 5G-H**). Addition of CsA hardly altered the rate or magnitude of Ca^2+^ release. Since deletion of MICS1 or LETM1 reduces KHE activity, we asked whether increasing KHE activity would restore Ca^2+^ release in MICS1KO mitochondria. Therefore, we repeated the previous experiment in the presence of nigericin, a highly selective ionophore catalyzing KHE, which did not restore Ca^2+^ efflux (**Figure 5I-J**). Thus, our results indicated that Na^+^-independent Ca^2+^ efflux requires MICS1 but not LETM1 or LETM1-mediated KHE activity.

### Thapsigargin-mobilized Ca^2+^ induces PTP opening in MICS1KO cells

The similar vigorous Ca^2+^ uptake by MICS1KO and -WT mitochondria but unequal Ca^2+^ release, unless alamethicin was used, raised the intriguing question of the fate of intramitochondrial Ca^2+^. To exclude the ER as a Ca^2+^ sink and deplete ER stores, we repeated Ca^2+^ uptake/release experiments using measurement media containing the SERCA pump inhibitor thapsigargin. MICS1WT mitochondria behaved as in the absence of thapsigargin, with identical rapid Ca^2+^ influx, RR-induced Ca^2+^ efflux and FCCP-induced release of total free matrix Ca^2+^ (**Figure 5K**). Ca^2+^ uptake was comparable in MICS1WT and MICS1KO#1, while significantly slowed in MICS1KO#2 (**Figure S2**). In contrast, MICS1KO#1 and MICS1KO#2 mitochondria, which were refractory to Ca^2+^ efflux in absence of thapsigargin, released RR-induced Ca^2+^ with rates 4-6 times higher than in MICS1WT. The levels of Ca^2+^ efflux seemed saturated, as they almost reached those of total Ca^2+^ release after FCCP addition, which in presence of thapsigargin were comparable to those of MICS1WT (**Figure 5K-L**). These drastic effects of thapsigargin on mitochondrial RR-induced Ca^2+^ efflux observed when MICS1 was deleted and NCLX inhibited, suggested stimulation of the CHE or opening of the PTP, which could both be caused by increased matrix Ca^2+^ load. Consistent with PTP opening (Beghi and Giussani, 2018), RR-induced Ca^2+^ release was accompanied by significant depolarization of MICS1KO but not MICS1WT mitochondria as indicated by the membrane potential dye TMRM (**Figure 5M-N**). To verify the PTP Ca^2+^-sensitivity and evaluate the total free Ca^2+^ load tolerated by MICS1KO mitochondria, we performed Ca^2+^ retention capacity (CRC) assays. MICS1WT mitochondria exposed to thapsigargin in presence of CGP37157 tolerated 5 Ca^2+^ pulses, corresponding to 25 μM Ca^2+^ before PTP opening **(Figure 5O)**. In contrast, MICS1KO#1 only tolerated 3 Ca^2+^ pulses, corresponding to 15 μM Ca^2+^ **(Figure 5P)**. PTP desensitization with CsA increased the retention capacity to a very similar extent in MICS1KO and MICS1WT mitochondria (**Figure 5O-P**). In the absence of CsA, the increased sensitivity to Ca^2+^-induced PTP opening of MICS1KO#1 required NCLX inhibition, as without addition of CGP37157 the retention capacity was the same for MICS1WT and MICS1KO#1 mitochondria with 20 μM Ca^2+^ **(Figure 5–figure supplement 2)**. The activation of PTP observed in **Figure 5K** indicated that thapsigargin increased the Ca^2+^ load in MICS1KO mitochondria. Likely, by inducing higher Ca^2+^ uptake rates or mobilizing an additional source of matrix Ca^2+^, and lowering the threshold for tolerated Ca^2+^, which would reduce Ca^2+^ buffering capacity. The CRC results confirmed the sensitization to Ca^2+^-induced PTP opening when NCLX is inhibited (Luongo et al., 2017). Consistent with a role of PTP opening in the large Ca^2+^ release observed in MICS1KO mitochondria, addition of CsA and ADP prevented excess Ca^2+^ release from MICS1KO mitochondria (**Figure 5Q-R**).

### Purified reconstituted MICS1 transports Ca^2+^

To assess the mechanism and selectivity of MICS1-dependent in cation transport we produced purified MICS1 for reconstitution studies. Codon optimized hMICS1 cDNA (**Figure 6–figure supplement 1A**) was cloned in pH6EX3 (Galluccio et al., 2013) and the recombinant construct was used to transform *E. coli* Rosetta cells. During the exponential phase of growth (OD ∼ 0.8-1), the temperature was set to 37 °C and 0.4 mM IPTG was added to induce synthesis of the protein. MICS1 was over-expressed in the insoluble fraction of the induced cell lysate after 2 hours of IPTG induction (**Figure 6–figure supplement 1B**). The protein was purified by Ni-chelating chromatography and reconstituted in proteoliposomes to assess in vitro Ca^2+^ transport activity assays using Calcium Green-5N as described Materials and Methods and illustrated in **Figure 6A**. The incorporation of MICS1 in proteoliposomes was verified by western blot analysis (**Figure 6B**). As shown in **Figure 6C-E,** reconstituted MICS1 mediated Ca^2+^ fluxes in a pH-dependent manner, with a maximum at pH 7.0 (**Figure 6D**) and inhibition of fluxes at pH 8.0 (**Figure 6E**). To further investigate the involvement of H^+^ in the transport cycle, we measured H^+^ flux using the pH sensitive dye pyranine (**Figure 6F**). Remarkably, alkalinization of the internal compartment of proteoliposomes detected by the increase in pyranine fluorescence indicated a H^+^ flux towards the external compartment induced by Ca^2+^ addition, i.e., concomitant to the inwardly directed Ca^2+^ flux (**Figure 6A**).

**Figure 6.**
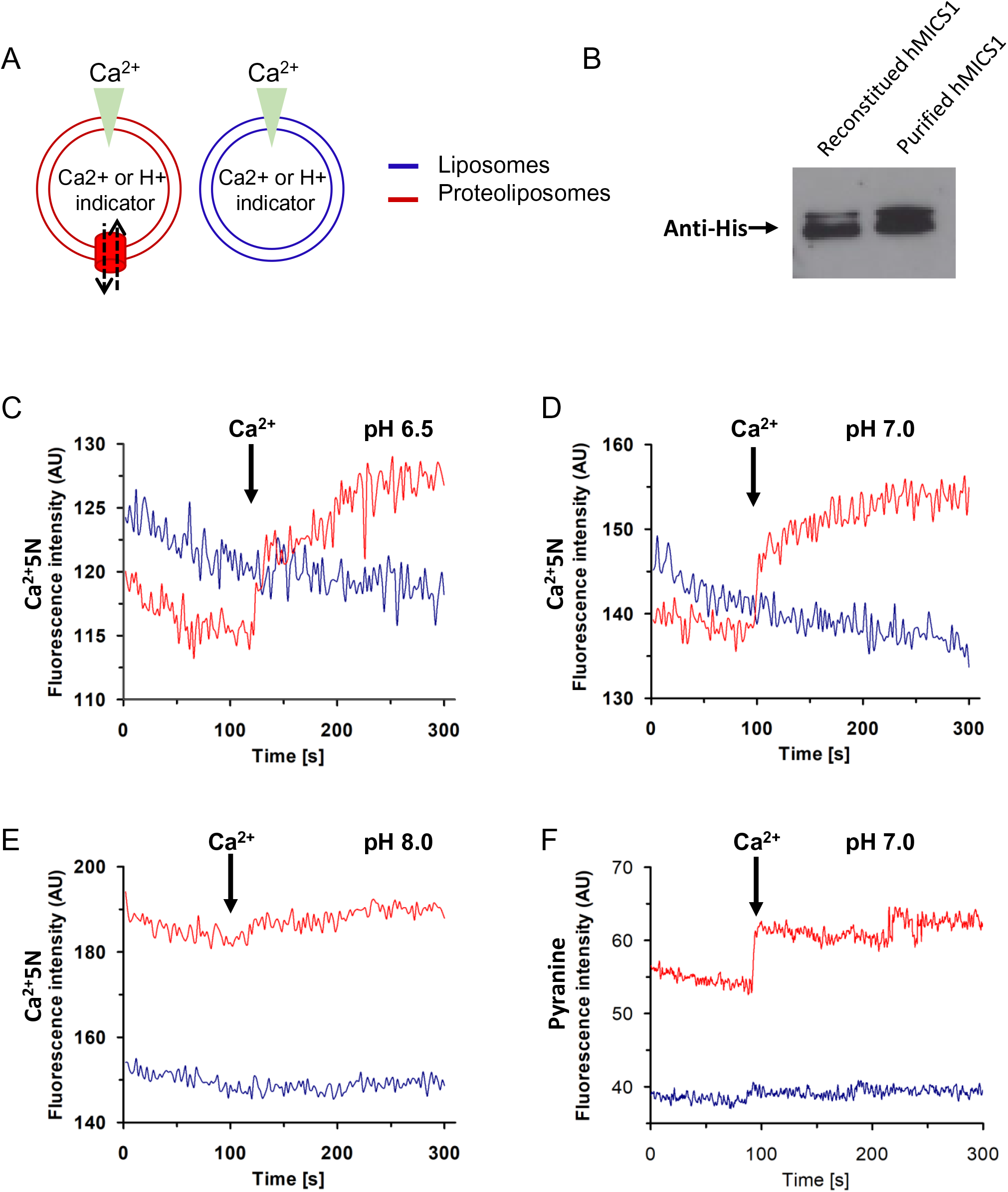
MICS1 proteoliposomes mediate Ca^2+^ and Ca^2+^-dependent H^+^ transport. **(A**) Sketch illustrating the reconstitution of hMICS1 (red) in proteoliposomes and the empty liposomes (blue) for transport measurements. **(B**) Western blot analysis of purified and reconstituted hMICS1 for evaluating the incorporation of hMICS1 into proteoliposomes prepared as described in materials and methods. Transport of Ca^2+^ **(C-E)** or H^+^ **(F)** by hMICS1 reconstituted in proteoliposomes. Purified hMICS1 was reconstituted in proteoliposomes containing 10 μM Calcium Green 5N at the pH indicated in the panels **(C-E)** or 20 μM pyranine at pH 7.0 **(F)**. After reconstitution, the fluorescence measurement was started by diluting 200 µL proteoliposomes (red trace) up to 3 mL with transport buffer prepared as described in materials and methods at the indicated pH **(C)** or at pH 7.0 **(D)**. After 100 sec, as indicated by the arrow, 7 mM Ca^2+^ was added to the sample and fluorescence change was recorded. As a control, the same measurement was performed diluting 200 µL liposomes (without incorporated protein, blue trace) up to 3 mL with the same transport buffer. The fluorescence intensity is indicated as Arbitrary Units (AU). Results are representative of three independent experiments. See also **Figure 6–figure supplement 1** for MICS1 optimization, induction and structure overview.

## Discussion

The role and selectivity of LETM1 as an ion transporter/channel has not been unequivocally assessed. The open questions remained whether it transports K^+^ or Ca^2+^, and whether it operates as an exchanger or rather as a component of the transport system. The work by Shao et al. showed that purified LETM1 oligomerizes into a high molecular weight complex of > 404 kDa, which forms a central cavity that undergoes pH-dependent conformational changes (Shao et al., 2016). In line with other reports, they proposed that LETM1 is a mitochondrial CHE (Doonan et al., 2014; Jiang et al., 2013). However, other studies clearly demonstrated that LETM1 plays a key role in mitochondrial K^+^ transport (Austin et al., 2017; Hashimi et al., 2013; Nowikovsky et al., 2012). The unresolved identity of the mitochondrial CHE and the controversy on LETM1 motivated us to further search for LETM1 interactors that could functionally cooperate with LETM1 in mitochondrial K^+^ and/or Ca^2+^ efflux.

To address the relatively low mitochondrial protein yield from mammalian cell cultures, we developed a miniaturized proteomic approach that was validated with the MCU interactome as a model. Among the most promising identified interactors of LETM1, we focused on MICS1.

Our study demonstrates that a complex containing LETM1 and MICS1 is involved in K^+^/H^+^ exchange in vivo, since decreased levels of both LETM1KD and MICS1KO led to a decrease of K^+^ transport. This effect is likely due either to reduced LETM1 levels, also in absence of MICS1, or to loss of protein interaction, a possibility that will be further explored in future analyses of MICS1 and LETM1 mutations affecting their physical interaction. Comparison of the roles of MICS1 and LETM1 in mitochondrial Ca^2+^ efflux clearly showed that LETM1, unlike MICS1, is not required for CHE activity. In contrast to LETM1, loss of MICS1 abrogated the function of CHE, which was restored by re-expression of MICS1. Independent of any interaction partner or protein complex, reconstituted MICS1 was able to transport Ca^2+^ across proteoliposomes in a pH-dependent manner and to drive Ca^2+^-dependent H^+^ transport. Thus, based on the consistency between cellular and cell-free activity of MICS1 in Na^+^-independent mitochondrial Ca^2+^ translocation, we have identified MICS1 as the long-sought mitochondrial CHE. Interestingly, MICS1 does not belong to any mitochondrial carrier family. The MICS1 structure predicted by AlphaFold (Jumper et al., 2021) shows a typical fold of membrane proteins with transport function with eight transmembrane segments and a long unresolved extra membrane domain (**Figure 6–figure supplement 1C**). MICS1 has no homologue in yeast *S. cerevisiae,* which lacks a mitochondrial Ca^2+^ uptake pathway.

As previously shown (Oka et al., 2008) and clearly confirmed here, loss of MICS1 causes changes in mitochondrial morphology. Our data additionally demonstrate that the morphological alterations are matched by reduced respiratory capacity that becomes evident with galactose as a substrate. The basis of this may reside in perturbation of Ca^2+^ homeostasis leading to excessive Ca^2+^ accumulation and possibly alterations of K^+^ homeostasis linked to secondary effects on LETM1. Thus, our findings link mitochondrial dysfunction to cation deregulation and provide a solid molecular framework for future studies.

The lack of a functional CHE also has severe implications on the permeability transition when Na^+^ dependent Ca^2+^ efflux is concomitantly blocked, as revealed by thapsigargin-induced hypersensitization of PTP opening. One reason that may explain why MICS1KO mitochondria are so sensitive to the PTP opening is reduced levels of Sirt3, which is responsible for deacetylation of CypD, a key PTP sensitizer (Sambri et al., 2020). The result is in accordance with the modulatory effect of thapsigargin on shifting the ratio between bound and free Ca^2+^ towards free Ca^2+^ (Korge and Weiss, 1999). The hypersensitivity of Ca^2+^-induced PTP opening also correlates with the observed cristae reorganization, OPA1 cleavage pattern and OMA1 activation, which could explain increased predisposition to cell death in MICS1KO cells exposed to thapsigargin.

In conclusion, the use of cell free and cell culture models has allowed us to demonstrate that MICS1 is the mitochondrial CHE. In view of the established involvement of LETM1 in both KHE and CHE activity, the identification of the LETM1 partner MICS1 is also a major step forward in resolving current controversies on their relative role in mitochondrial Ca^2+^ and K^+^ homeostasis. Indeed, we have demonstrated that MICS1 is a necessary part of the KHE machinery and its interaction with LETM1 fulfills a physiological role in the cell and in maintaining Ca^2+^ balance. Further investigation of LETM1 and MICS1 interaction partners will shed further light on the regulatory mechanism maintaining mitochondrial ion balance.

## Material and Methods

### Reagents

All reagents used in this study were from Sigma Aldrich, unless otherwise indicated.

Antibiotics: normocin, blasticidin, hygromycin, puromycin and doxycycline were from Invivogen (San Diego, CA). Restriction endonucleases and specific reagents for cloning, Pierce BCA protein assay kit, Glutaraldehyde, lead citrate, propylene oxide and osmium tetroxide were from Merck (Darmstadt, Germany), ProtA/G agarose and DMEM (#41966-029) from Thermo Fisher Scientific, NativeMark™ #LC0725) NativePAGE™ 3-12 % Bis-Tris Protein, # BN1001), Turbofect, Ca2+ Green 5N, and MitoTracker™ Green FM (#M7514) from Invitrogen, Streptactin beads from IBA-lifesciences. Bradford was from BioRad, Proteinase inhibitor from Roche (Basel, Switzerland), C12E8 from TCI Europe, TMRM from Molecular Probes, Glycid ether 100 from Serva (Heidelberg, Germany). Fetal Bovine Serum (FBS), and pen/strep were from Gibco. Mycoplasma test kit was from MycoAlert Lonza kit. The working concentration of Ruthenium red was calculated with Lambert-Beer law, A=533 nm, I = 1 cm, Ɛ = 65000.

### Antibodies used in this study

LETM1 (Abnova, #H00003954), 1:1000, LETM1 (Santa Cruz Biotechnology, #sc-163013), 1:1000, LETM1 C-terminal region (Aviva, Systems Biology, #OAAB12878), 1:1000, MICS1 (Abcam, #ab106754), 1:1000, MICS1 (Aviva systems biology, #OAAF06415), 1:1000, OPA1 (BD Biosciences, #612606), 1:1000, SIRT3 (Cell Signaling Technology, #5490), 1:1000, DRP1 (Santa Cruz Biotechnology, #sc-271583), 1:1000, HSP60 (Santa Cruz Biotechnology, #sc-1052), 1:1000, TOM20 (Cell Signaling Technology, #42406), 1:1000, TOM40 (Santa Cruz Biotechnology, #sc-365467), 1:1000, OMA1 (Santa Cruz Biotechnology, #sc-515788), 1:1000, Prohibitin (Abcam, #ab210082), 1:1000, β-Actin (Invitrogen, #MA5-11869), 1:1000, PolyHistidine-Peroxidase (Sigma Aldrich, #a7058), 1:10000, Goat-α-mouse (Jackson ImmunoResearch, #115-035-003) 1:5000, Rabbit-α-goat (Jackson ImmunoResearch, #305-035-003) 1:5000, Goat-α-rabbit (Jackson ImmunoResearch, #111-035-144) 1:5000.

### Cell culture

HEK293 Flp-In T-Rex (Invitrogen), HeLa (Austin et al., 2017) and HEK293 (ATCC) cells were maintained in DMEM supplemented with FBS (10% v/v), and penicillin/streptomycin (pen/strep) (1%). Cells were cultured in an incubator set to 37 °C and 5% CO_2_ and splitted when reaching confluency of ∼70-90%, and regularly tested for mycoplasma.

### Generation of knockdown and knockout cells

All shRNA constructs for MICS1 and LETM1 were obtained from Origene Technologies (Rockville, MD). Primers were from Microsynth, Balgach, Switzerland. HeLa scramble and LETM1KD cells were described in (40). HEK293 scramble and LETM1 knockdown cells were generated using the short hairpin constructs from (Austin et al, 2017) in HEK293-Flp-In T-Rex cells. MICS1KD cells were generated using the human shRNA plasmid kit for MICS1 (Origene, TR315671B) with the shRNA construct #1 (GGTCTTGGAGCATTCTGCTACTATGGCTT) and construct #2 (GCCATAGCAATCAGCAGAACGCCTGTTCT and GGTCCTCTTCTCATCAGAGCTGCATGGTA) used for stable KD cell lines. Cells were transfected with Turbofect according to the manufacturer instructions; 48 hrs post transfection the media was changed to selection media containing puromycin (2 µg/mL). Puromycin-resistant cell populations were maintained in growth media supplemented with puromycin (1 µg/mL). MICS1KO cells were generated by the Protein Technologies Facility at Vienna BioCenter Core Facilities (VBCF), member of the Vienna BioCenter (VBC), Austria. 4 gRNAs targeting GHITM were designed using CRISPOR tool (crispor.tefor.net). gRNAs were selected primarily on the criterium of their specificity (at least 3 mismatches with at least one in the seed region to any off-target) and on predicted activity according to Doench score. Guide 1: CCAAAACAAGAATTGGGATC (targeting exon 3), guide 2: GCATTGTGCTACTATGGCTT (targeting exon 4), guide 3: CAGCCATTGATTCTTCGTGA (targeting exon 2) and guide 4: GGCTCCTCTGACAATATTA (targeting exon 7). Targeting sequences were introduced into pX459 Cas9-p2A-puro plasmid (Addgene 48139) via BbsI cloning. Plasmids (3 µg) were introduced into HEK 293 cells (1×10^6^) by electroporation with Neon electroporator (Thermo Fisher Scientific) according to the manufacturers protocol. 24 hrs post electroporation cells were selected with puromycin (4 µg/ml) and 72 hrs later collected and lysed for genotyping. Editing efficiency was confirmed with TIDE algorithm (https://tide.deskgen.com/) based on chromatogram analysis with WT HEK293 PCR product used as a reference. Guide 2 (GCATTGTGCTACTATGGCTT) was selected for performing the KO in HEK and HeLa cells based on its highest activity (59.7%) and cloned into an in-house template vector p31 containing T7 promoter followed by BbsI cloning sites, optimized gRNA scaffold and DraI restriction site used for template linearization. Resulting gRNA transcription was performed with HiScribe T7 High Yield RNA Synthesis Kit (NEB) according to the manufacturer’s protocol and gRNA was purified and verified for concentration and RNA integrity. 12 µg of gRNA pre-mixed with 5 µg Cas9 protein (2×NLS) in Cas9 buffer (20 mM HEPES pH 7.5, 150 mM KCl, 0.5 mM DTT, 0.1 mM EDTA) were used for electroporation of 70-80 % confluent cells. Electroporated cells were cultured in DMEM supplemented with 10 % FCS and L-Gln. Normocin was added after approximately 2 hrs, and after 24 hrs genotyping was performed to confirm editing.

### Mitochondria isolation

Frozen cell pellets were thawed and resuspended in isolation media (Austin et al., 2017) containing 1.7 mM Proteinase inhibitor cocktail. Cells were homogenized on ice with 12 strokes at 1600 rpm with a Yellowline OST basic homogenizer and mitochondria isolated by differential centrifugation according to (Frezza et al., 2007).

### Molecular cloning

#### LETM1 for mass spectrometry experiments

Expression constructs for LETM1-SH were PCR amplified from pVT-U LETM1 (Nowikovsky et al., 2004) and subcloned into the pTO-SII-HA-GW vector which was a kind gift from M. Gstaiger (ETH, Zurich). Subcloning was done by Gateway cloning (Invitrogen, Carlsbad, CA). Plasmid (pTO-SII-HA-GW GFP) expressing N-terminal tagged GFP with Strep-HA tag was a kind gift from A. Bergthaler (CeMM, Vienna). Primers: attB LETM1 forward 5’GGGGACAAGTTTGTACAAAAAAGCAGGCTAGACTGCCATGGCGTCCAT3’, attB LETM1 reverse 5’GGGGACCACTTTGTACAAGAAAGCTGGGTTGCTCTTCACCTCTGCGAC3’.

#### MICS1 for rescue experiments

The human MICS1 cDNA was amplified by reverse-transcriptase PCR using the forward primer 5’AAGCTTGACCATGTTGGCTGCAAGG3’, and the reverse primer with in frame Flag sequence 5’GTCTCTCGAGTTACTTGTCATCGTCATCCTTGTAATCTTTCTTTCTGTTGCCTCC3’ and cloned into the pcDNA3 Plasmid (Sigma Aldrich) using the restriction sites *HindIII* and *Xho1*.

#### MICS1 for proteoliposomes

Codon optimization of the human MICS1 sequence (UniProtKB: Q9H3K2; GenPept accession no. NP_055209.2) was designed using Genscript and increased the Codon Adaptation Index (CAI) from 0.32 to 0.97. The codon optimized cDNA encoding for human MICS1 protein was sub-cloned from pUC57 by double digestion and inserted between *Hind*III and *Xho*I restriction sites of the pH6EX3 expression vector. The resulting recombinant plasmid encodes a 6His-tagged fusion protein corresponding to the hMICS1 carrying the extra N-terminal sequence MSPIHHHHHHLVPRGSEA.

### Generation of stable LETM1-StrepHA or MICS1-Flag expressing cell lines

Cells were transfected with Turbofect according to the manufacturer instructions; 48 hrs post transfection the media was changed to selection media as according to the resistance marker of the plasmid. The concentration of selection antibiotics as listed: hygromycin (260 µg/mL), blasticidin S (38 µg/mL). After a resistant population of cells was established, cells were maintained in growth media containing: hygromycin (100 µg/mL), blasticidin S (15 µg/mL). For stable MICS1-Flag cells G418 (1 mg/mL) was used in media devoid of FBS and pen/strep.

### AP-MS sample preparation

N-terminally tagged GFP or LETM1 inducible HEK293 Flp-In T-Rex were generated as outlined above. Protein expression was induced with doxycycline (1µg/mL) for 24 hrs in standard culture media. Cells were lysed and the bait protein purified by affinity purification (AP) as described (Rudashevskaya et al., 2013). Affinity purification from mitochondria was performed as (Rudashevskaya et al., 2013) with modification. Crudely isolated mitochondria were lysed using 6-aminocarprotic acid with protease inhibitors and n-Dodecyl β-D-maltoside (2% w/v) and vortexed for 30 min at 4 °C. Lysates were cleared at 15000 ×g, 4 °C for 15 min and the supernatant was quantified by Bradford assay with BSA as standard. Protein complexes were purified from 2 mg crude mitochondrial input with Streptactin (IBA, Göttingen, Germany) beads. Washing steps were performed in scaled volume of AP buffer, thrice with detergent, twice without and then eluted with biotin (Alfa-Aesar, Ward Hill, MA). Protein complexes were reduced, alkylated and digested with trypsin as described (Rudashevskaya et al., 2013). Peptides were desalted and concentrated by reversed-phase tips (Rappsilber et al., 2007) and reconstituted in formic acid (5%) for LC-MS analysis.

### Reversed-phase LC-MS data analysis and data filtering

All liquid chromatography mass spectrometry experiments were performed on an Aglient 1200 HPLC nanoflow system coupled to a linear trap quadrupole (LTQ) Orbitrap Velos mass spectrometer (ThermoFisher Scientific). Raw data were matched to peptides and proteins using Mascot and Phenyx, with a false discovery rate of 1% at the protein level. CRAPome and SAINT analysis were applied to all AP-MS data. GFP pulldowns were used as controls together with publically-available CRAPome data that used similar sample preparation and MS methods and instrumentation. Common contaminants and proteins with a frequency greater than or equal to 0.1 in the CRAPome database were excluded. Proteins with a SAINT score greater than 0.97 were identified as high confidence interactors.

### Co-immunoprecipitation

Cells were washed with PBS and harvested in coIP buffer (150 mM NaCl, 50 mM Tris, 2 mM EDTA, 1% IGEPAL C360, and protease inhibitor tablet without EDTA. Cells lysates were vortexed, cleared and quantified as described above. Lysates (500 µg or 1 mg) were then incubated overnight with Streptactin beads or primary antibody as indicated (10 µg). Primary antibody samples were then incubated for 1 hr at RT with ProtA/G agarose. Beads were then washed 3 times with coIP buffer then 2 times with PBS and eluted with 1X Laemmli buffer for SDS-PAGE and immunoblotting.

### Western blotting: SDS and BN PAGE

SDS PAGE and immunoblotting were performed as in (Austin et al., 2017). Bradford or BCA assays were performed according to the manufactureŕs protocol and blots were quantified using the BioRad Image Lab (v6.1.0) software. For BNGE, isolated mitochondria were solubilized with a final concentration of 1% digitonin for 15 min on ice, centrifuged at 27000 ×g for 30 min in a Beckman Optima^TM^ ultracentrifuge and the supernatant (corresponding to 5 µg) with G-250 Sample Additive (0.5 µl) was separated using precasted gels (NativePAGE™ 3-12 % Bis-Tris Protein). Unstained Protein Standard NativeMark™ served as a marker. Protein complexes were transferred onto PVDF membranes overnight using wet blotting at 30 V.

### Proliferation assay

Cells were seeded counted manually every 24 hrs by trypan blue exclusion. At least three independent counts were performed on each sample. Cell numbers were plotted and data shown as means ± SD.

### Light scattering assays

Light scattering experiments were adapted from previous protocols (Austin et al., 2017) Briefly, mitochondria were isolated from HEK293 and HeLa cells as described in (Frezza et al., 2007) and resuspended in isolation media (200 mM Sucrose, 10 mM Mops-TRIS, 1 mM EGTA-TRIS, pH:7,4). Antimycin A (5 μM) was used at RT to depolarize mitochondria and A123187 (1 μM) and EDTA (10 μM) to deplete matrix magnesium. Light scattering assays were conducted in a photometric 96 well plate reader (Varioscan) at RT; KOAc media (180 μl), as described in (Austin et al., 2017) was injected to 200 μg mitochondria (total volume 200 μl) and absorbance was detected at OD_540 nm_. Quinine (0.5 mM) served to inhibit the KHE. The swelling rate was quantified by one phase decay on raw swelling data as shown, K value as rate constant.

### Ca^2+^ uptake/release assays

Cells (7 x 10^6^) were permeabilized with digitonin (1.25 %) in 400 µl permeabilization media PM1 (KCl (130 mM), Mops-Tris pH 7.4 (10 mM) EGTA-Tris (1 mM), KPi pH.7.4 (1 mM). Permeabilization was stopped (immediately after 80-90% of the cells had become permeable to trypan blue) in 600 µl PM2 (KCl (130 mM), Mops-Tris pH 7.4 (10 mM), EGTA-Tris (10 µM), KPi pH.7.4 (1 mM), and resuspended in measurement media contained sucrose (250 mM), MOPS-Tris (10 mM), EGTA-Tris (10 µM), KPi 7.4 (1 mM), sodium succinate (5 mM) rotenone (2 µM), sodium succinate (5 mM) to energize CII and rotenone (2 µM) to block CI. CGP37157 (2 µM) served to inhibit NCLX, and when indicated thapsigargin (1 µM) to block SERCA. Calcium Green-5N (0.24 µM) was used to record extramitochondrial Ca^2+^, TMRM (0.33 µM) to measure the membrane potential. A bolus of CaCl_2_ (10 µM) was applied to initiate Ca^2+^ uptake. MCU was inhibited by addition of RR (0.2 µM), which induced Ca^2+^ release. FCCP (2 µM) or alamethicin (2.5 µM) was added to induce the maximal release of total Ca^2+^ at the end of the measurement. The LS55 spectrofluorometer 211 (Perkin Elmer) was used with the following parameters: Ca^2+^ green-5N: λ_ex_= 505 nm, λ_em_ = 530 nm, slit width: Ex-2.5 nm, Em-2.5 nm; TMRM: λ_ex_= 546 nm, λ_em_ = 590 nm, slit width: 2.5 nm.

### Calcium Retention Capacity experiment

The measurements were performed in media as described for Ca^2+^ uptake release assay, containing Calcium Green-5N and when indicated thapsigargin (1 µM) CsA (1 µM) and/ or CGP37157 (1 µM). CaCl_2_ pulses (5 µM) were added sequentially until the opening of PTP occurred. Measurements were performed using the LS55 spectrofluorometer 211 (Perkin Elmer) with the same parameters as for Ca^2+^ uptake release.

### Seahorse Mito Stress assay

Extracellular flux analyses were performed with the Agilent Seahorse XF24 Extracellular flux analyser as outlined in (Wilfinger et al., 2016), with minor modifications to inhibitor concentration, oligomycin (0.5 μM) and FCCP (0.2 μM). Carbon source is indicated in the figure legends, either (glucose 25 mM) or galactose (10 mM), all media were supplemented with sodium pyruvate (1 mM).

### Transmission electron microscopy

Cells were fixed in glutaraldehyde (5%) phosphate buffer (0.1 M) (Sigma–Aldrich, Vienna, Austria), pH 7.2, at 4 °C for 2 hrs. Subsequently, samples were post-fixed in 1% osmium tetroxide in the same buffer at 4 °C for 1 hr. After dehydration in an alcohol gradient series and propylene oxide, the tissue samples were embedded in glycid ether 100. Ultrathin sections were cut on a Leica ultramicrotome (Leica Ultracut S, Vienna, Austria), stained with uranyl acetate and lead citrate and examined with a Zeiss TEM 900 electron microscope (Carl Zeiss, Oberkochen, Germany) operated at 80 kV.

### Live cell imaging

For confocal microscopy 5 x 10^4^ cells/well were seeded onto poly-L-lysine coated μ-Slide 8 well plates (Ibidi, #80826). The next day mitochondria were loaded with MitoTracker™ Green FM (50 nM) for 30 minutes and then changed to fresh medium before they were monitored under 5 % CO_2_ at 37 °C using a LSM880 microscope with Plan-Apochromat 63x/1.40 Oil DIC M27 lens. MTG was excited at a wavelength of 488 nm and images were processed in Adobe Photoshop CS2.

### Over-expression, purification and reconstitution in proteoliposomes of MICS1 for Ca^2+^ transport assays *Expression of MICS1 protein*

To produce the 6His-MICS1 recombinant protein, *E. coli* Rosetta cells (Novagen) were transformed with the pH6EX3-hMICS1 construct. Selection of transformed colonies was performed on LB-agar plates added with ampicillin (100 µg/mL) and chloramphenicol (34 µg/mL). A colony was inoculated and cultured overnight at 37 °C under rotary shaking (160 rpm). The day after, the culture was diluted 1:20 in fresh medium added with the specific antibiotics. When the optical density measured at OD_600 nm_ wavelength was 0.8-1, different IPTG concentrations (from 0.1 to 1 mM) were tested to induce protein expression except for one aliquot, grown in absence of inducer (negative control). The cultures were continued for up to 6 hours at 28 °C or 37 °C at 160 rpm. Every two hours, aliquots were collected and centrifuged at 3000 ×g, and at 4 °C for 10 minutes; the pellets were stored at −20 °C. A bacterial pellet aliquot, after thawing, was dissolved in a resuspension buffer (20 mM Hepes Tris, 200 mM NaCl pH 7.5) added with protease inhibitor cocktail according to manufacturer instructions. The bacterial suspensions were sonicated in an ice bath for 10 minutes (pulse of 1 second on, and 1 second off) at 40 Watt, using a Vibracell VCX-130 sonifier. The insoluble cell fractions were analyzed by SDS-PAGE and western blotting.

#### Purification of hMICS1

hMICS1, over-expressed in *E. coli*, was purified by Ni-chelating chromatography. In brief, the insoluble fraction of bacterial cell lysates was firstly washed with a buffer containing Tris-HCl pH 8.0 (0.1 M). After centrifugation step (12000 ×g for 5 min at 4 °C), pellet was resuspended with 100 mM 1,4-dithioerythritol (DTE) and then solubilized with a buffer containing urea (3.5 M), sarkosyl (0.8%), NaCl (100 mM), glycerol (5%), Tris HCl pH 8.0 (10 mM). After solubilization, the sample was centrifuged at 12000 ×g for 10 min at 4 °C and the supernatant was applied onto a column filled with 2 mL His select nickel affinity gel (0.5 cm diameter, 2.5 cm height) pre-conditioned with 8 mL of a buffer containing sarkosyl (0.1%), NaCl (200 mM), glycerol (10%), Tris HCl pH 8.0 (20 mM). Then, 5 mL of a buffer containing Tris HCl pH 8.0 (20 mM), glycerol (10%), NaCl (200 mM), n-Dodecyl β-D-maltoside (0.1%) and DTE (5 mM) was used to wash the column removing unbound proteins. In order to increase the purity of the recovered MICS1, another washing step was performed using 3 mL of the same above-described buffer added with 10 mM imidazole. Finally, MICS1 was eluted in 5 fractions of 1 mL, using the same above-described buffer added with 50 mM imidazole. The purified protein was eluted in a peak of 2.5 mL. The eluted protein was subjected to a buffer change for imidazole and Na^+^ removal, using a PD-10 column pre-conditioned with a desalt buffer composed of Tris HCl pH 8.0 (20 mM), glycerol (10%), n-Dodecyl β-D-maltoside (0.1%) and DTE (10 mM): 2.5 mL of the purified protein were loaded onto the PD10 column and collected in 3.5 mL of desalt buffer.

#### Reconstitution in proteoliposomes of the purified hMICS1

The desalted hMICS1 was reconstituted by removing detergent from mixed micelles of detergent, protein and phospholipids using the batch wise method previously described for other membrane proteins (Cosco et al., 2020), with some modifications to increase the protein/phospholipid ratio required for fluorometric measurements (Scalise et al., 2020). The initial mixture contained: 25 μg of purified protein, 50 μL of 10% C_12_E_8_, 50 μL of 10% egg yolk phospholipids (w/v) in the form of liposomes prepared as previously described (Scalise et al., 2018), 20 mM Tris HCl pH 7.0, except where differently indicated, 10 μM of Calcium Green-5N or 20 μM pyranine, in a final volume of 700 μL. The detergent was removed by incubating the reconstitution mixture with 0.5 g of the hydrophobic resin Amberlite XAD-4 for 40 min under rotatory stirring at room temperature.

#### Cation transport measurements by spectrofluorometric assays

The Ca^2+^ flux or the intraliposomal pH changes were monitored by measuring the fluorescence emission of Calcium Green-5N or pyranine, respectively included inside the proteoliposomes. After reconstitution, 600 µL of proteoliposomes was passed through a Sephadex G-75 column, pre-equilibrated with Tris HCl pH 7.0 (20 mM), except where differently indicated. Then, 200 µL proteoliposomes were diluted in 3 mL of the same buffer and incubated for 10 min in the dark prior to measurements. To start the transport assay, CaCl_2_ (7 mM) buffered at pH 7.0, except where differently indicated, was added to proteoliposomes; the uptake of Ca^2+^ or the efflux of H^+^ was measured as an increase of Calcium Green-5N or pyranine fluorescence, respectively. As a control, the same measurements were performed using liposomes, i.e., vesicles without reconstituted hMICS1. The measurements were performed in the fluorescence spectrometer (LS55) from Perkin Elmer under rotatory stirring. The fluorescence was measured following time drive acquisition protocol with λ excitation=506 nm and λ emission=532nm (slit 5/5) for Calcium Green-5N and λ excitation=450 nm and λ emission=520nm (slit 5/5) for pyranine.

### Statistical analysis

All statistical analyses were done in GraphPad (La Jolla, CA) Prism v6 for Windows. Bar graphs were generated with GraphPad Prism. Tests and individual *p* values as indicated in figure legends. The data are presented as mean ± SD unless specified.

## Acknowledgement

We thank Dr Martha Giacomello for constructive discussion and experimental support and Dr Paolo Bernardi for critical reading of the manuscript. This work was supported by the Doc fellowship from the Austrian Academy of Science ÖAW to SA and the Austrian Science Funds FWF research project grants P-314717 and P-29077 to KN.

## Author contributions

KN conceived the study and supervised the experimental work. SA performed the interactome, co-immunoprecipitation, bioenergetics, generated LETM1 and MICS1 knockdowns and analysed the data. RM performed K^+^, Ca^2+^ flux, CRC and delta psi measurements and data analysis, SM conducted immunofluorescence, cell biological and protein analysis, MS and MG generated *E. coli* strains, purified and reconstituted proteins and performed cell-free flux measurements, CP performed BNGE and TB LETM1 SDS-PAGE. KB designed and supervised mass spectrometry analysis, KP and DV participated in method development and running of LC-MS instrumentation, ND conducted TEM, CI designed the cell-free study, KN, SA and CI wrote the manuscript.

## Supplemental Figure titles and legends

**Figure 1- figure supplement 1.**
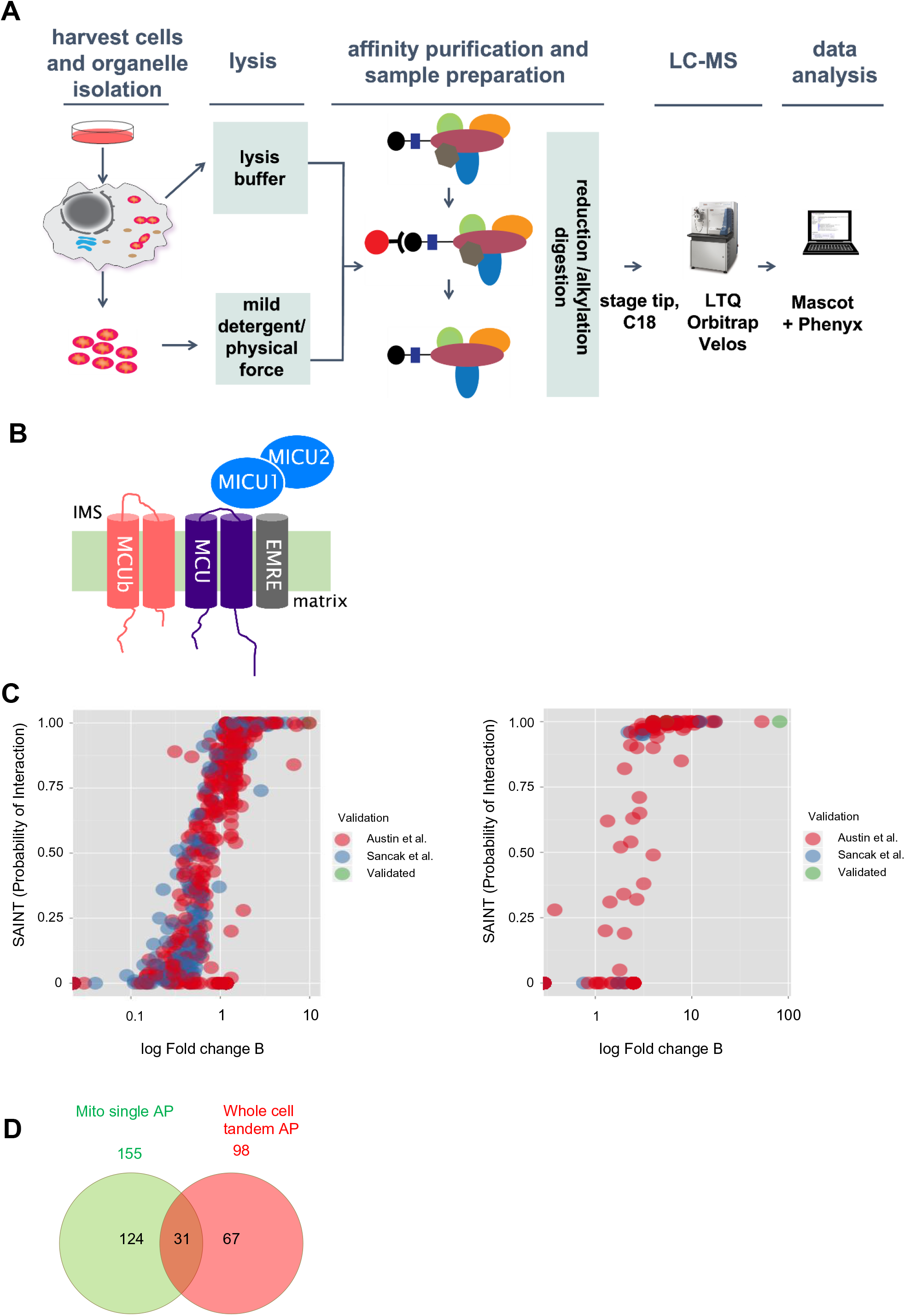
AP-MS experiments. (A) Scheme illustrating workflow for minaturized AP-MS experiments, left to right: whole cells or isolated mitochondria are lysed or solubilized respectively. The cell/mitochondrial lysates are used for affinity purification (AP) using the StrepHA tag found on the bait protein. Eluates of the AP and control experiments are reduced, alkylated, and digested by trypsin. Peptides are purified on a C18 stage tip and then run on a LTQ Orbitrap Velos. Protein identifications were made by internal tools using MASCOT and Phenyx and removal of non-specific interactors done using the CRAPome. (B) Mitochondrial calcium uniporter was selected as a model protein, the functional complex consist of the 5 proteins above (MCU, MCUb, MICU1, MICU2, EMRE). Note that an additional tissue-specific tertiary interaction partner (MICU3) **(2-4)**, is only expressed at very low levels in HEK293 cells (Diego De Stefani, personal communication). Illustration adapted from Sancak et al. (C) Proteins identified by AP-MS were scored for probability of interaction using SAINT score and fold change B using raw data and the Crapome (left), proteins identified in Austin et al (red) and Sancak et al (blue). Both GFP & Crapome controls of similar experimental setup were used for analysis to filter nonspecific interactors (right), proteins identified in Austin et al (red), Sancak et al (blue) and validated interaction partners found in both (green). (D) Common affinity purification contaminants were eliminated from GFP and MCU analysis by removing proteins that had greater than 5 average spectral counts in the Crapome Database. (E) Venn diagram illustrating the number of proteins identified in mitochondrial single streptavidin AP (156 proteins) or whole cell tandem AP (98 proteins). While 31 proteins were found in common with both approaches, with 124 proteins being unique to the mitochondrial single streptavidin AP and 67 being unique to the whole cell tandem AP.

**Figure 4-figure supplement 1.**
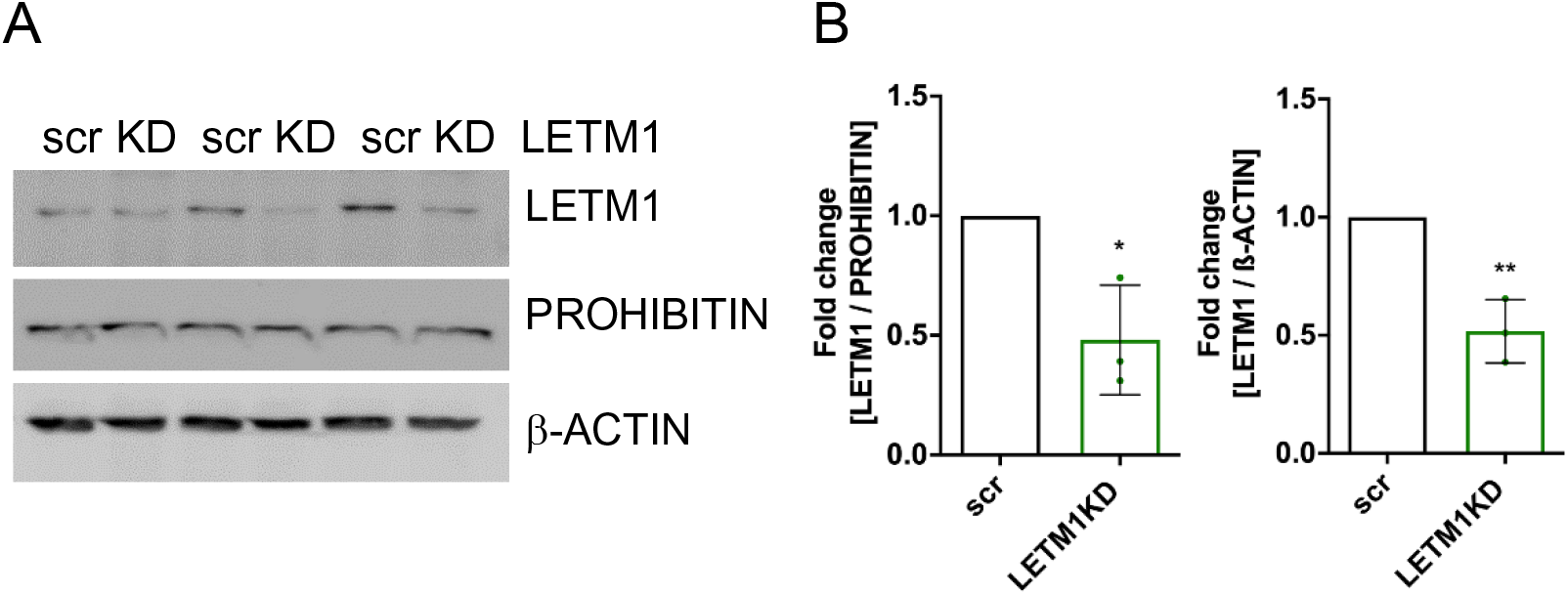
Downregulation of LETM1. (A) Western blot of HEK293 LETM1 scramble (scr) and LETM1KD (KD) used in Fig 4D-E. PROHIBITIN and β-ACTIN served as mitochondrial and total cellular loading control, respectively. (B) Statistics: unpaired two-sided t-test, *p <0.05, **p <0.01.

**Figure 5-figure supplement 1.**
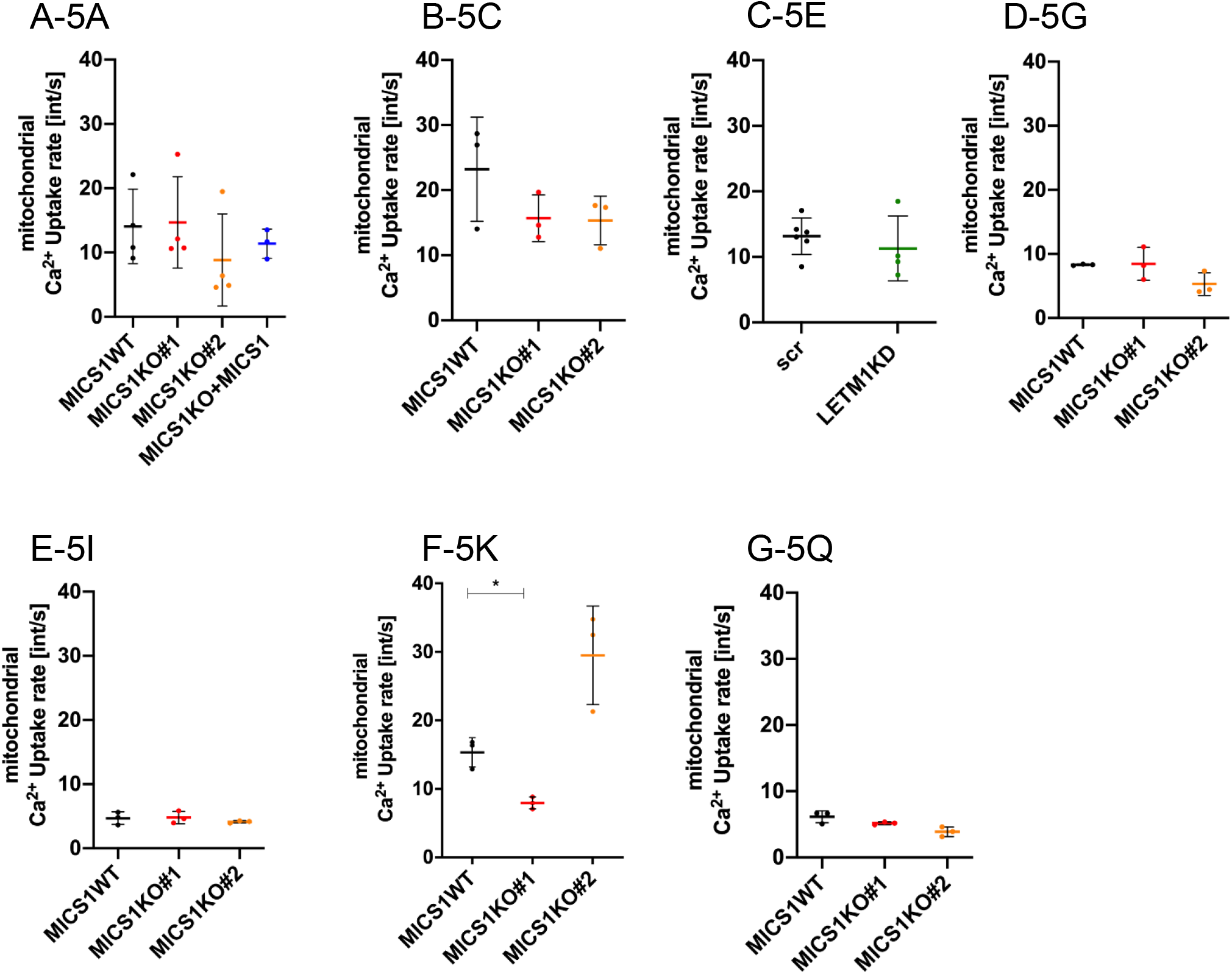
Mitochondrial Ca^2+^ uptake rates. The decay rate was calculated as exponential decrease of the Ca^2+^ 5N Signals using the equation: Y=(Y0 - Plateau)*exp(-K*X) + Plateau, Rate of Decay/sec: K*Y0; X: Time (sec), Y: Starts at Y0 (Ca^2+^ peak) and decays with one phase down to Plateau; K: Rate constant equal to the reciprocal of the X axis units. Quantification of the Ca^2+^ uptake rates recorded in Fig 5A (A), Fig 5C (B), Fig 5E (C), Fig 5G (D), Fig 5I (E), Fig 5K (F) and Fig 5P (G). Calculations were done using the GraphPad software and statistical analysis using Brown-Forsythe and Welch ANOVA, n=3, statistical analysis *p<0.05 for (F).

**Figure 5-figure supplement 2.**
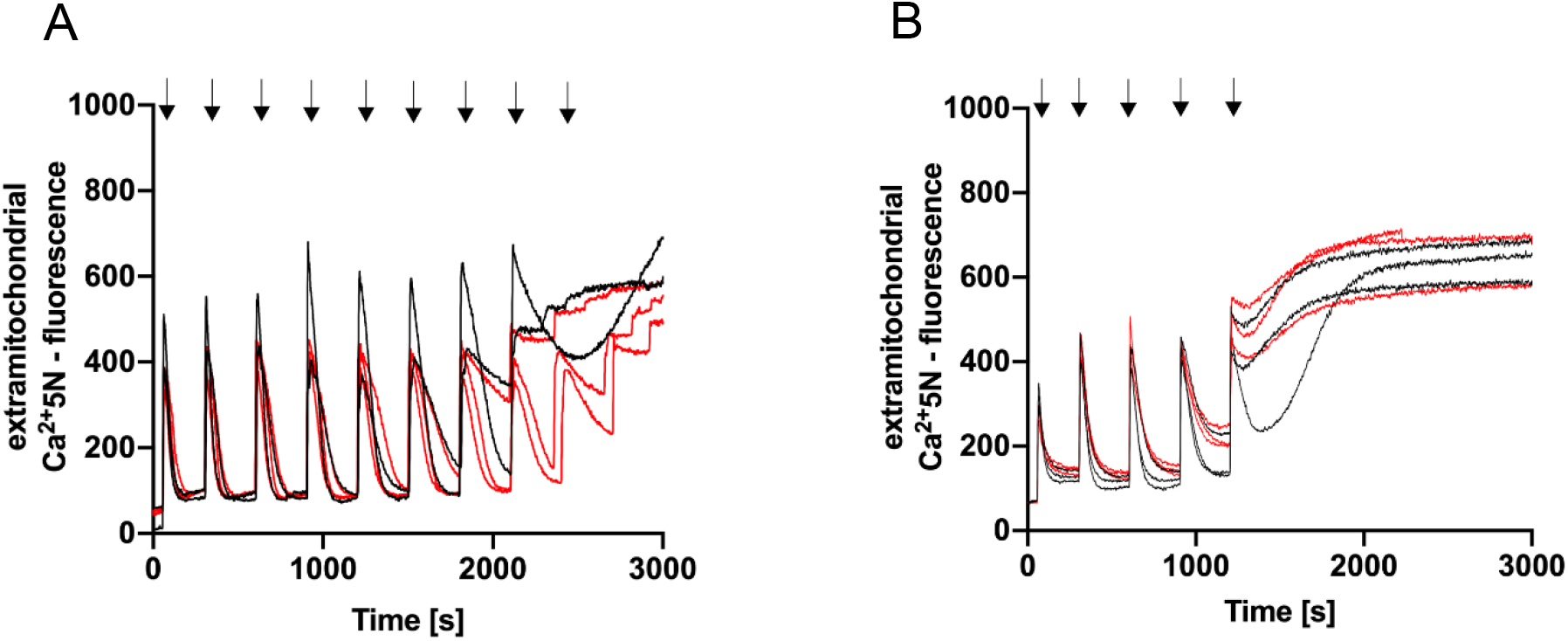
CRC in absence of CGP37157. Permeabilized HEK293 MICS1 WT (black trace) and MICS1 KO (red trace) cells exposed (A) or not (B) to CsA were subjected to sequential Ca^2+^ bolus of 5 µM Ca^2+^ while NCLX was not inhibited and fluorescence intensity was recorded.

**Figure 6-figure supplement 1.**
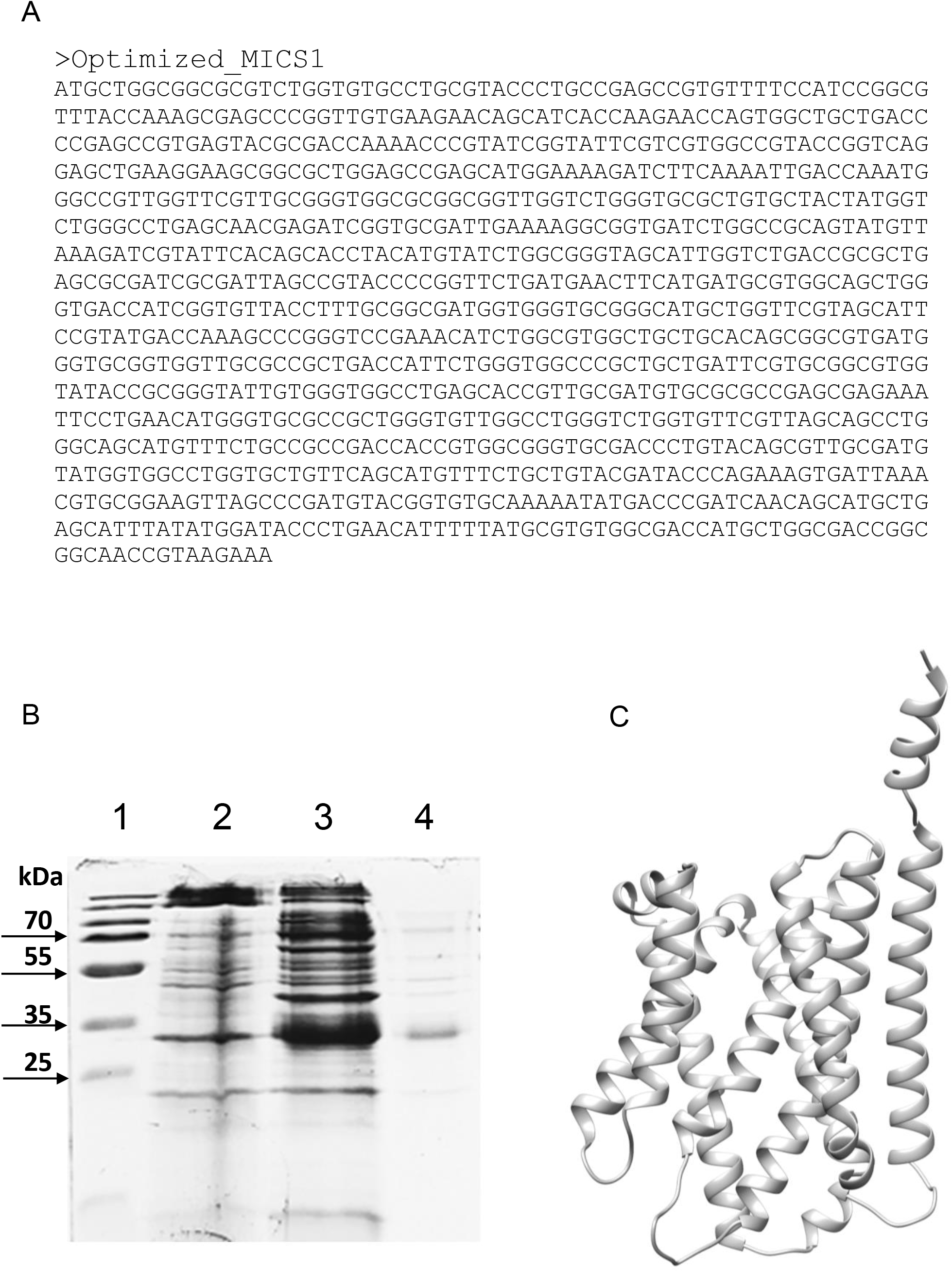
MICS1 optimization, induction and structure overview. A) Nucleotide sequence of the codon optimized sequence of MICS1 (B) MICS1 expression and purification. Lane 1: page ruler prestained plus marker; lane 2: insoluble fraction of not induced cell lysate (negative control); lane 3: insoluble fraction of cell lysate after 2 hours of 0.4 mM IPTG induction at 37 °C; lane 4: purified MICS1 protein. (C) Ribbon representation of the hMICS1 protein (Q9H3K2). PDB file retrieved from AlphaFhold at https://alphafold.ebi.ac.uk/entry/Q9H3K2. The unstructured loop containing the first 55 amino acids has been removed.

